# Biomechanical effects of adding an articulating toe joint to a passive foot prosthesis for incline and decline walking

**DOI:** 10.1101/2023.12.05.570262

**Authors:** Rachel H. Teater, Karl E. Zelik, Kirsty A. McDonald

## Abstract

Walking on sloped surfaces is challenging for many lower limb prosthesis users, in part due to the limited ankle range of motion provided by typical prosthetic ankle-foot devices. Adding a toe joint could potentially benefit users by providing an additional degree of flexibility to adapt to sloped surfaces, but this remains untested. The objective of this study was to characterize the effect of a prosthesis with an articulating toe joint on the preferences and gait biomechanics of individuals with unilateral below-knee limb loss walking on slopes. Nine active prosthesis users walked on an instrumented treadmill at a +5° incline and −5° decline while wearing an experimental foot prosthesis with two configurations: a Flexible toe joint and a Locked-out toe joint. Three participants preferred the Flexible toe joint over the Locked-out toe joint for incline and decline walking. Eight of nine participants went on to participate in a biomechanical data collection. The Flexible toe joint decreased prosthesis Push-off work by 2 J during both incline and decline walking (p=0.008). During incline walking, prosthetic limb knee flexion at toe-off was 3° greater in the Flexible configuration compared to the Locked (p=0.008). Overall, these results indicate that adding a toe joint to a passive foot prosthesis has relatively small effects on joint kinematics and kinetics during sloped walking. This study is part of a larger body of work that also assessed the impact of a prosthetic toe joint for level and uneven terrain walking and stair ascent/descent. Collectively, toe joints do not appear to substantially or consistently alter lower limb mechanics for active unilateral below-knee prosthesis users. Our findings also demonstrate that user preference for passive prosthetic technology may be both subject-specific and task-specific. Future work could investigate the inter-individual preferences and potential benefits of a prosthetic toe joint for lower-mobility individuals.

## Introduction

Typically-able-bodied individuals adapt to walking on sloped surfaces by altering their gait mechanics. For example, relative to walking on level ground, they exhibit increased magnitudes of ankle dorsiflexion on both inclines and declines (1,2). For walking uphill, the biological ankle also provides additional positive work to move the center-of-mass up the slope against gravity. During downhill walking, the lower limbs mostly absorb energy and generate less positive work at the joints as the center-of-mass is lowered (3–6).

Unlike able-bodied individuals, people with lower limb loss cannot easily alter their prosthetic ankle’s flexion or power generation to accommodate a slope (7,8). As a result, many lower limb prosthesis users (LLPUs) struggle to comfortably navigate sloped terrains. A study that surveyed over 300 LLPUs found that only 52% of users were able to walk on uneven ground such as sloped surfaces without assistance (9). LLPUs have also been observed to have reduced walking speed and cadence when walking on slopes (7,8) and increased metabolic cost on inclines (10) compared to individuals without limb loss. These challenges associated with sloped walking can have substantial impacts on a person’s physical activity level, quality-of-life, and societal participation.

Most LLPUs use a passive ankle-foot prosthesis that has a fixed ankle set-point angle and rigid foot segment (11). This results in limited ankle range of motion and power generation from the device, which likely contribute to gait compensations observed during sloped walking (8,12–14). For example, individuals with below-knee limb loss often exhibit altered knee mechanics compared to individuals without limb loss when walking on slopes (8,15). During incline walking, users have reduced knee flexion in early stance accompanied by reduced dorsiflexion provided by the prosthetic ankle (8,15). During decline walking, the reported increase in prosthetic limb knee flexion during late stance is thought to compensate for the lack of prosthetic ankle range of motion when lowering one’s center-of-mass (8,15,16).

Device interventions that aim to improve the ability of LLPUs to walk on sloped surfaces have often focused on adapting the behavior of the prosthetic ankle joint. Passive, hydraulic, microprocessor, and fully powered devices have all been investigated to determine if they can improve the ability of users to walk on slopes (10,12–19). Some studies have found a benefit in their approach to modulating ankle behavior to improve incline and decline walking, but most have reported mixed or conflicting results. For example, a study investigating a microprocessor device that adjusts the set-point of the ankle joint during sloped walking reported an increase in ankle range of motion and prosthetic limb knee flexion during incline walking, but for declines the added plantarflexion did not show a measurable benefit and biomechanical results conflicted with participant feedback on the device (15).

An alternative or complementary approach to adjusting device ankle behavior during sloped walking may be to incorporate a passive toe joint into a prosthetic foot. Biological metatarsophalangeal (i.e., toe) joint articulation plays an important role in locomotion and prior experimental and simulation research suggests that altering toe joint dynamics can impact gait mechanics and locomotor economy (20–27). Incorporating a toe joint into a prosthetic foot could benefit LLPUs during locomotion in general, or for specific locomotor tasks. We have previously investigated level ground walking with a toe joint added to a passive ankle-foot prosthesis (28). During this study, we found that a Flexible toe joint decreased the amount of Push-off power provided by the prosthesis compared to a Locked-out toe joint configuration but observed no significant difference in rate of oxygen consumption or the biomechanics of other lower limb joints. Four of nine participants preferred the Flexible configuration for level ground walking.

For walking on slopes, an articulating toe joint could provide LLPUs with an additional degree of compliance to help compensate for the lack of prosthetic ankle flexion. For walking uphill, a prosthesis with a flexible toe joint may increase the ability of the device to conform to the sloped surface, but potentially has the drawback of providing less Push-off power compared to a prosthetic foot of the same design without a toe joint. For walking downhill, a prosthesis with a toe joint could potentially aid LLPUs by providing flexibility in the device during late stance to support lowering their center-of-mass. The potential reduction in prosthetic Push-off power may be inconsequential or potentially beneficial when walking downhill as there is less need to generate positive power with the lower limbs and specifically the ankle joint (3–6). However, the impact of incorporating a flexible toe joint into a prothesis for sloped walking has never been tested before. The potential gait adaptations or benefits and preferences of LLPUs are unknown.

Therefore, the objective of this study was to characterize the preferences and gait biomechanics of unilateral below-knee prosthesis users walking on an incline and decline wearing a prosthetic foot in two configurations: a Flexible and Locked-out toe joint. Specifically, we evaluated user preference, spatiotemporal variables, prosthesis Push-off work, and prosthetic limb knee kinematics to determine the impact of a Flexible toe joint for sloped walking. We expected positive prosthesis Push-off work to be reduced for the Flexible configuration compared to the Locked configuration for both incline and decline walking. We expected LLPUs to prefer the Locked configuration for incline walking due to the increased amount of positive work (from the ankle and at the center-of-mass level) that is necessary to ascend ramps (3–6). For decline walking, we expected users to prefer the Flexible configuration as it provides an additional degree of flexibility during late stance to assist LLPUs in lowering their center-of-mass as they descend ramps. We also predicted that early stance kinematics of the prosthetic limb would not be influenced by the addition of a toe joint, but late stance kinematics would be affected. This was evaluated by quantifying prosthetic limb knee angle at initial contact and toe-off.

## Methods

### Participants

Nine below-knee prosthesis users (male, age: 40.7 ± 10.5 years, mass: 95.0 ± 12.9 kg, height: 1.84 ± 0.05 m; Table 1) participated in a multi-day research protocol (28–30), which included incline and decline walking. Participant recruitment for this study started on November 1st, 2018 and concluded on September 15th, 2019. Each participant provided written informed consent, according to Vanderbilt University’s Institutional Review Board procedures. Participant 7 was not able to attend the session when incline and decline biomechanical walking data were collected due to scheduling constraints and therefore only eight participants are included in the biomechanical analyses. All participants were a Medicare Functional Classification Level of K4, except one (K3). All were able to walk without a mobility aid and were at least six months post amputation surgery at the time of data collection.

**Table 1.**
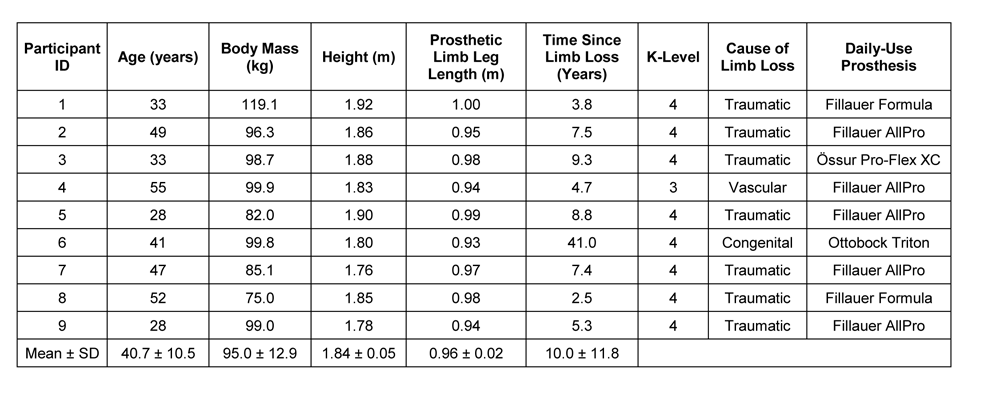
Individual and mean (± standard deviation) participant demographics.

### Experimental prosthesis

The experimental device was a commercial prosthetic foot (Balance Foot J, Össur, Categories 25, 27 and 28) that was modified to function in two configurations (Fig 1). The first configuration was a Flexible toe joint that was accomplished by attaching a truncated foot keel to a toe segment with sheets of spring steel allowing toe joint flexion during loading (Fig 1A). The stiffness of the toe joint was 0.34 N m degree^−1^. The Locked toe joint configuration was accomplished by connecting the foot and toe segments with a rigid aluminum block that prevented toe joint flexion to effectively create a solid foot keel without a toe joint (Fig 1B). During data collection, participants wore a cosmesis over the experimental prosthesis, with no shoe.

**Fig 1.**
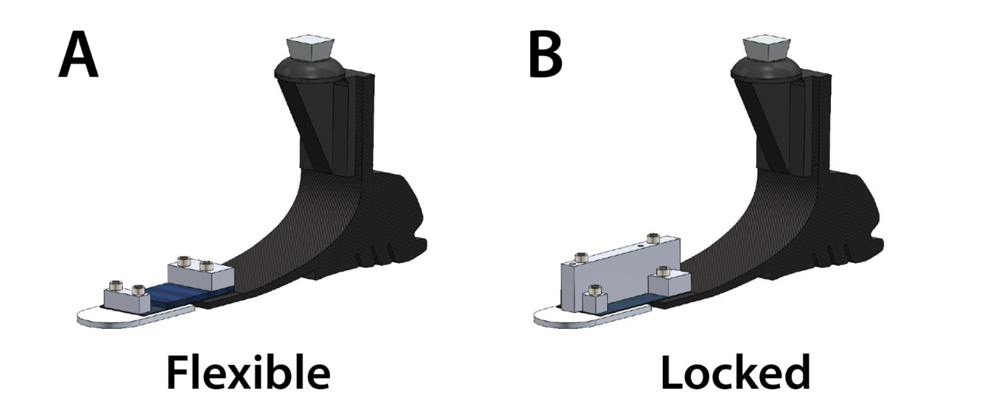
Experimental prosthetic foot. A commercial prosthetic foot (Össur Balance Foot J) modified to function in two configurations: (A) a Flexible toe joint configuration made using sheets of spring steel and (B) a Locked toe joint configuration accomplished by securing an aluminum block across the joint to prevent flexion.

### Training protocol

The full research protocol involved four sessions: two training and two testing. During the first training session, participants familiarized themselves with six locomotor tasks (walking over level, incline, decline, and uneven terrain surfaces, and ascending/descending stairs) while wearing their prescribed prosthesis. They were then fitted with the experimental prosthesis in a randomized starting configuration (Flexible or Locked). A certified prosthetist conducted the alignment of the experimental prosthesis, which was maintained for all training and testing sessions. Over the two training sessions, participants acclimated to the six previously mentioned tasks in both the Flexible and Locked configurations. The order of the two configurations for training and testing was randomized across participants. At the end of the second training session, participants ranked their satisfaction with their familiarization on the experimental prosthesis in each configuration for all locomotor tasks on a scale of 1-10. All participants reported a 9 or 10 across tasks and configurations, indicating they felt comfortable and acclimated to walking on the experimental prosthesis. At the conclusion of the second training session, participants were also asked to report their preference for Foot One or Foot Two for each locomotor task. ‘Foot One’ and ‘Foot Two’ referred to the randomized configuration order (Locked then Flexible or Flexible then Locked) for each participant.

### Testing protocol and data collection

Data collection for incline and decline walking in each toe joint configuration was performed on the fourth day of the protocol (second testing session). Participants had 40-42 reflective markers affixed to their lower limbs and the experimental prosthesis (28). Kinematics were simultaneously collected at 200 Hz (10-camera system, Vicon, Oxford, UK) with ground reaction forces (GRFs) collected at 1,000 Hz while participants walked at 1 m s^−1^ on a dual-belt force-measuring treadmill (Bertec, Columbus, OH, USA) at a +5° and −5° slope. Speed and slopes were chosen based on existing literature examining prosthesis user locomotion (10,18,31). The duration of the walking trial for both the Flexible and Locked configuration was approximately 90 s, with the middle 60 s of data collected for analysis.

### Outcome Metrics

In this study, we provide time-series angle, moment, and power data for all lower limb joints during both incline and decline walking. However, we chose a select number of key outcome metrics to evaluate the impact of incorporating a toe joint into a passive ankle-foot prosthesis. Peak prosthesis toe joint angle was computed to confirm that our experimental prosthesis functioned as intended in both configurations. We computed positive prosthesis Push-off work to evaluate the hypothesized difference in energy provided by the device in the two configurations. We also characterized prosthetic limb knee angle at initial contact and toe-off to determine whether adding a toe joint impacts the altered knee flexion angles typically observed for LLPUs walking on slopes compared to individuals without limb loss (8,15,32).

### Data analysis

Motion capture and GRF data were filtered with a fourth order, low-pass Butterworth filter with a cutoff frequency of 8 and 15 Hz, respectively. Spatiotemporal results (stride length and time; stance and swing time for each limb) were computed using a 20 N threshold for vertical GRF (to identify initial contact and toe-off of each limb) and the position of the posterior calcaneus marker on each limb. Sagittal plane joint angles and net moments, and net joint powers (six-degree-of-freedom) and center-of-mass dynamics (33) were computed via inverse dynamics using Visual3D (C-motion, Germantown, USA) and custom MATLAB code (MathWorks, Natick, USA). Prosthesis power and work were calculated using the distal segment power method (34,35). Prosthesis work was computed for the Push-off phase of gait using the positive range of prosthesis power during late stance. Data for each participant and condition were further processed in MATLAB and divided into strides then normalized to 100% of the stride cycle. An average of 40 strides per trial were included for analysis with included strides determined by clean force plate contacts (e.g., strides where the foot contacted two force plates at once were omitted).

Outcome metrics were then non-dimensionalized to account for differences in participant size. Stride length was non-dimensionalized by leg length (*L*). Stride time, stance time, and swing time were non-dimensionalized by √*L/g*, where *g* is acceleration due to gravity (36,37). Moments and work were non-dimensionalized by *MgL* where *M* is body mass. Power was non-dimensionalized by *Mg*√*gL* (36,37). Average non-dimensionalization constants were 0.96 m (length), 0.31 s (time), 893.5 N m and J (moment and work), and 2853.6 W (power). Some outcomes were re-dimensionalized using these constants for reporting purposes.

### Statistical analysis

Group results for relevant outcome metrics are reported as mean ± standard deviation. Data were screened for normality via a Shapiro-Wilk test. Following this, a series of paired-samples t-tests (normal distribution) or Wilcoxon signed-rank tests (non-normal distribution) were applied to detect differences in spatiotemporal variables, peak toe joint angle, prosthesis Push-off work, prosthetic limb knee angle at initial contact, and prosthetic limb knee angle at toe-off between the Flexible and Locked configuration trials. Holm-Bonferroni corrections were applied to account for familywise error rates across the groups for spatiotemporal and principal kinematic and kinetic variables. Incline and decline data were considered separate families. Adjusted alpha levels for each variable during incline walking are listed in parentheses: stride length (p=0.017), stride time (p=0.010), prosthetic limb stance time (p=0.013), prosthetic limb swing time (p=0.025), intact limb stance time (p=0.008), intact limb swing time (p=0.05), peak toe joint angle (p=0.013), prosthesis Push-off work (p=0.018), prosthetic limb knee angle at initial contact (p=0.050), and prosthetic limb knee angle at toe-off (p=0.025). Adjusted alpha levels for each variable during decline walking are listed here: stride length (p=0.017), stride time (p=0.050), prosthetic limb stance time (p=0.010), prosthetic limb swing time (p=0.013), intact limb stance time (p=0.025), intact limb swing time (p=0.008), peak toe joint angle (p=0.0125), prosthesis Push-off work (p=0.018), prosthetic limb knee angle at initial contact (p=0.05), and prosthetic limb knee angle at toe-off (p=0.025). Statistical analyses were conducted in MATLAB and Excel (Microsoft, Redmond, USA).

## Results

Group-level average curves for lower limb kinematics and kinetics divided by limb and slope are provided in Figs 2-5. Most results were normally distributed. Non-normally distributed variables for incline walking included prosthetic limb swing time, intact limb swing time, prosthesis Push-off work, and prosthetic limb knee angle at initial contact. Non-normally distributed variables for decline walking included stride time, intact limb stance time, and prosthesis Push-off work.

**Fig 2.**
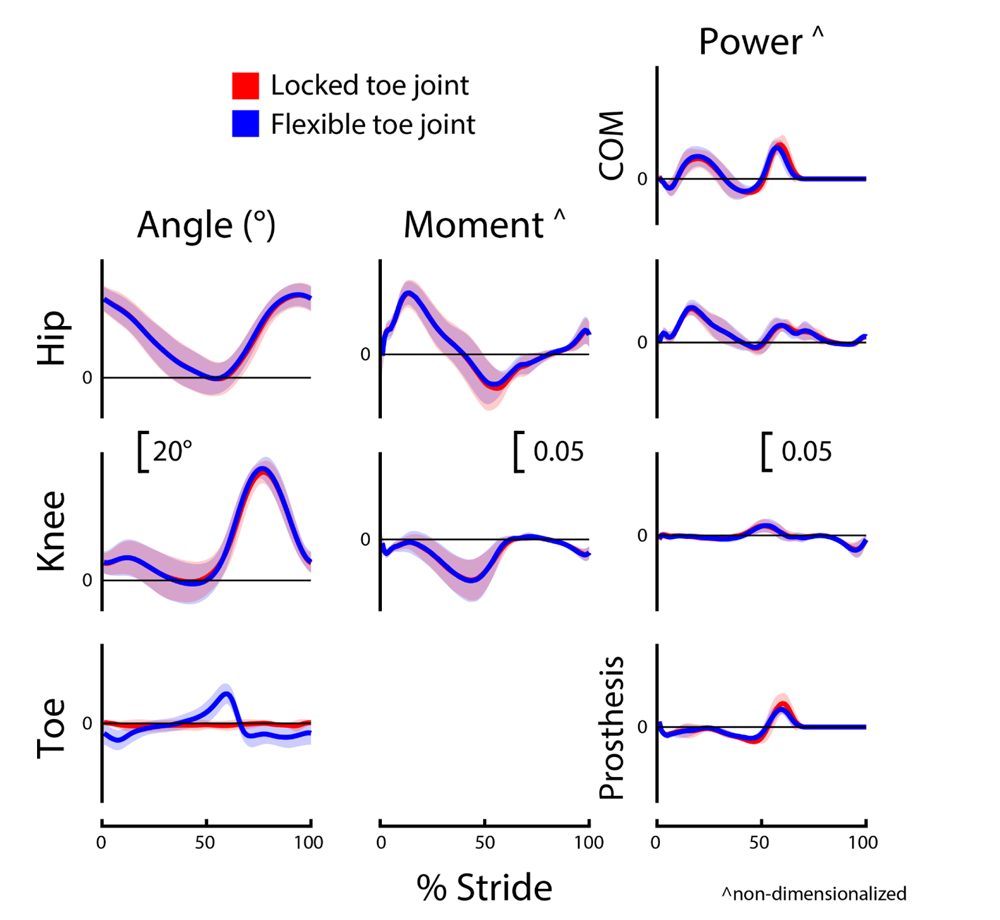
Prosthetic limb biomechanics during incline (+5°) walking. Prosthetic limb joint, center-of-mass (COM), and prosthesis dynamics for participants (*N=8*) using a passive prosthetic foot with a Flexible (blue) and Locked (red) toe joint configuration. Data were cropped into strides using prosthetic limb heel strikes. Data are presented as mean ± standard deviation (shaded regions). Moment and power data are presented as dimensionless values. Using group mean re-dimensionalization constants, 0.05 corresponds to 0.47 N m kg^−1^ for moments and 1.50 W kg^−1^ for powers.

**Fig 3.**
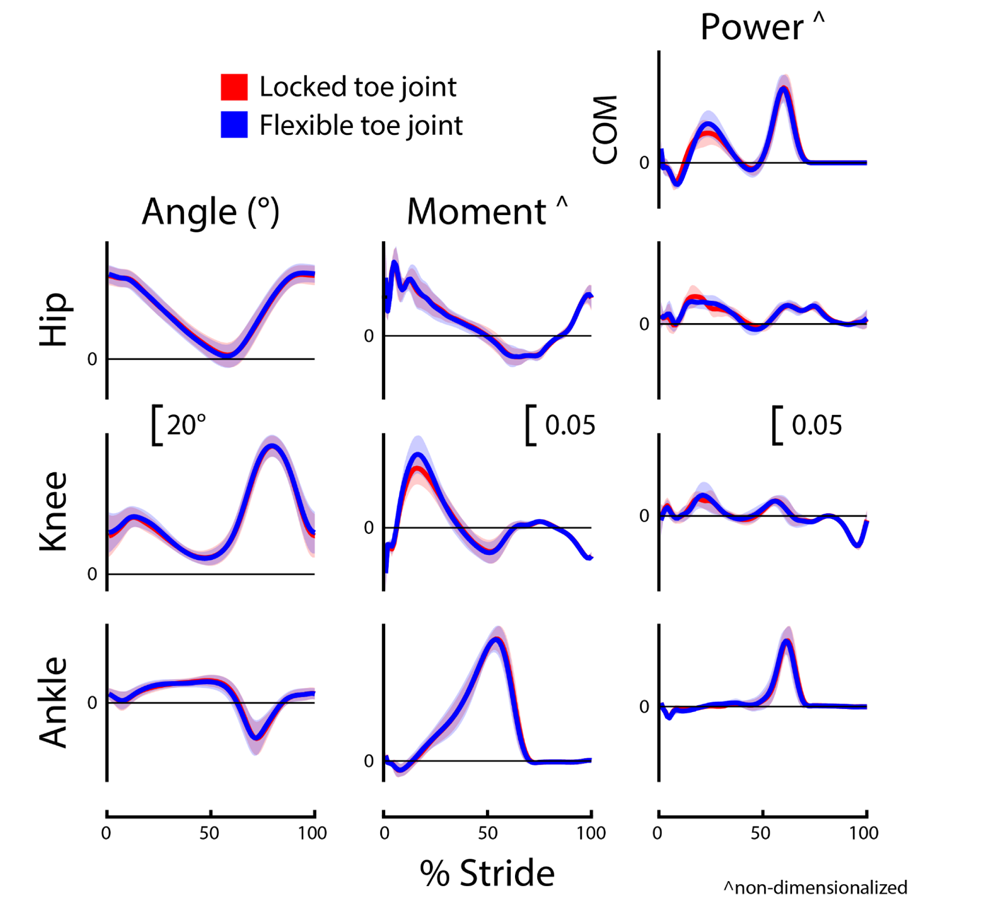
Intact limb biomechanics during incline (+5°) walking. Intact (non-prosthetic) limb joint and center-of-mass (COM) dynamics for participants (N=8) using a passive prosthetic foot with a Flexible (blue) and Locked (red) toe joint configuration. Data were cropped into strides using intact limb heel strikes. Data are presented as mean ± standard deviation (shaded regions). Moment and power data are presented as dimensionless values. Using group mean re-dimensionalization constants, 0.05 corresponds to 0.47 N m kg^−1^ for moments and 1.50 W kg^−1^ for powers.

**Fig 4.**
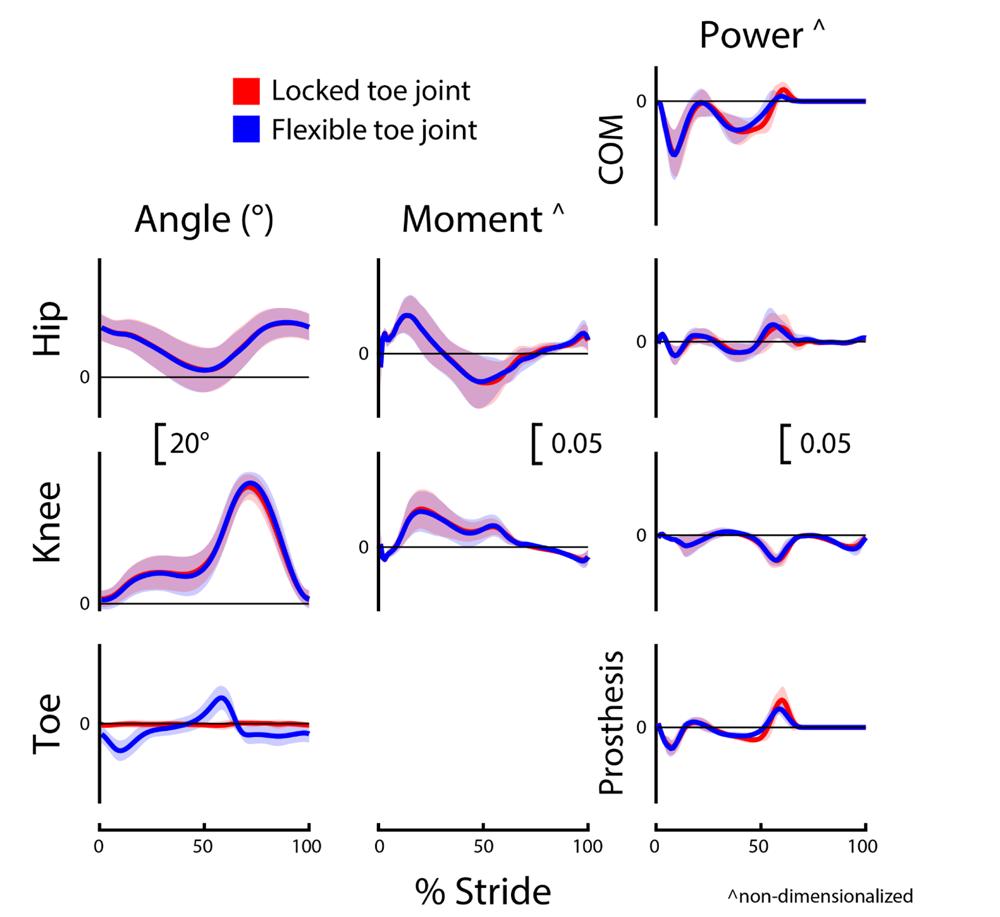
Prosthetic limb biomechanics during decline (-5°) walking. Prosthetic limb joint, center-of-mass (COM), and prosthesis dynamics for participants (*N=8*) using a passive prosthetic foot with a Flexible (blue) and Locked (red) toe joint configuration. Data were cropped into strides using prosthetic limb heel strikes. Data are presented as mean ± standard deviation (shaded regions). Moment and power data are presented as dimensionless values. Using group mean re-dimensionalization constants, 0.05 corresponds to 0.47 N m kg^−1^ for moments and 1.50 W kg^−1^ for powers.

**Fig 5.**
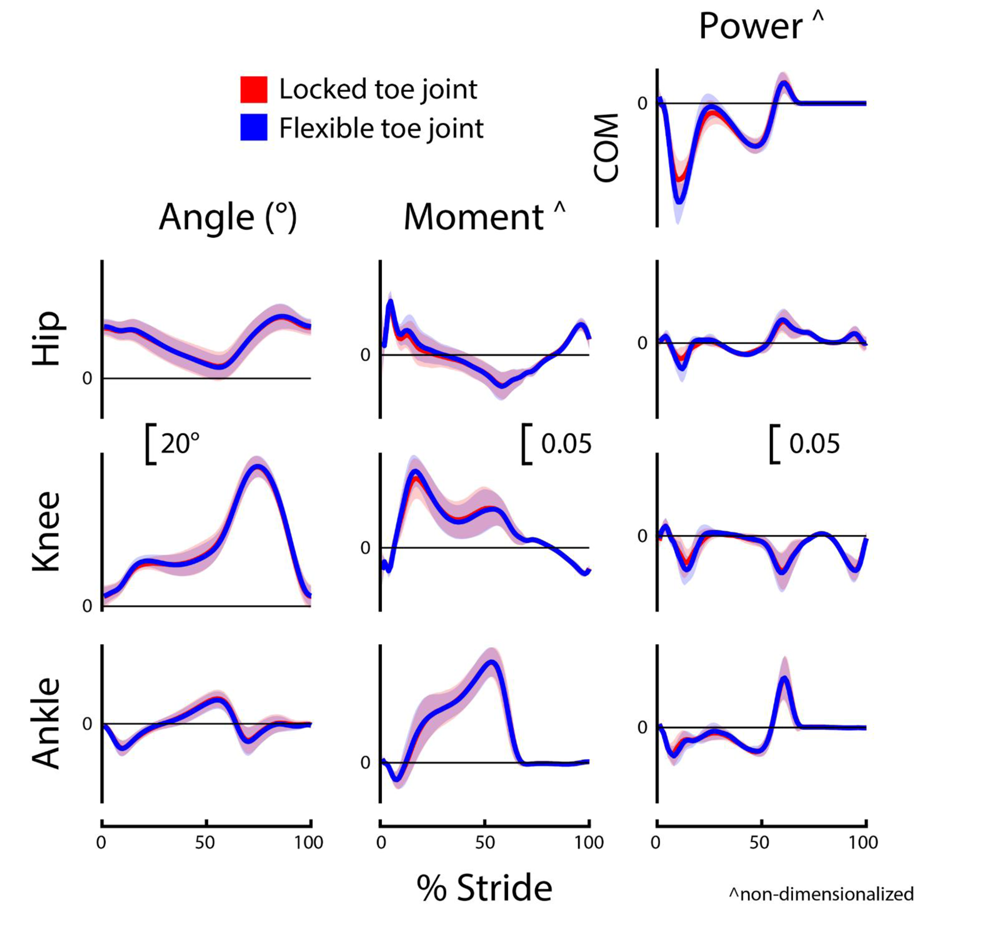
Intact limb biomechanics during decline (-5°) walking. Intact (non-prosthetic) limb joint and center-of-mass (COM) dynamics for participants (*N=8*) using a passive prosthetic foot with a Flexible (blue) and Locked (red) toe joint configuration. Data were cropped into strides using intact limb heel strikes. Data are presented as mean ± standard deviation (shaded regions). Moment and power data are presented as dimensionless values. Using group mean re-dimensionalization constants, 0.05 corresponds to 0.47 N m kg^−1^ for moments, and 1.50 W kg^−1^ for powers.

### User preference

Participant preference was divided between the Flexible and Locked configuration for both incline and decline walking. For incline walking, three of nine participants (Participants 1, 3, and 4) preferred the Flexible toe joint while the remaining six preferred the Locked toe joint. For decline walking, three of nine again preferred the Flexible configuration while the remaining six preferred the Locked, but it was a slightly different group of three participants (Participants 1, 2, and 3).

### Spatiotemporal

Spatiotemporal results are presented in Table 2. During incline walking, there was only a significant difference between the Flexible (0.77 s) and Locked (0.79 s) configurations for intact limb stance time (p=0.008). During decline walking there were no significant differences in spatiotemporal variables between configurations (p>0.48).

**Table 2.** Mean (± standard deviation) re-dimensionalized spatiotemporal variables.

|  |  | Stride Length<br>(m) | Stride Time<br>(s) | Prosthetic Limb |  | Intact Limb |  |
| --- | --- | --- | --- | --- | --- | --- | --- |
|  |  |  |  | Stance Time (s) | Swing Time (s) | Stance Time (s) | Swing Time (s) |
| Incline<br>(+5°) | Flexible | 1.13 ± 0.09 | 1.13 ± 0.09 | 0.72 ± 0.07 | 0.41 ± 0.04 | 0.77 ± 0.06* | 0.36 ± 0.04 |
| | Locked | 1.15 ± 0.09 | 1.15 ± 0.09 | 0.74 ± 0.08 | 0.41 ± 0.08 | 0.79 ± 0.06* | 0.37 ± 0.03 |
| Decline<br>(-5°) | Flexible | 1.03 ± 0.07 | 1.03 ± 0.07 | 0.65 ± 0.06 | 0.38 ± 0.03 | 0.68 ± 0.05 | 0.35 ± 0.04 |
| | Locked | 1.03 ± 0.07 | 1.03 ± 0.07 | 0.64 ± 0.06 | 0.38 ± 0.03 | 0.68 ± 0.05 | 0.35 ± 0.04 |

\*Significant difference detected between the Flexible and Locked configurations.

### Toe joint angle

During incline walking, the peak toe joint angle for the Flexible configuration (18.0 ± 3.7°) was significantly greater than the Locked configuration (1.7 ± 1.3°; p<0.001; Table 3; Fig 2). There was also a significant difference in peak toe joint angle during decline walking between the Flexible configuration (16.7 ± 4.6°) and Locked configuration (1.5 ± 1.5°; p<0.001; Table 3; Fig 4).

**Table 3.**
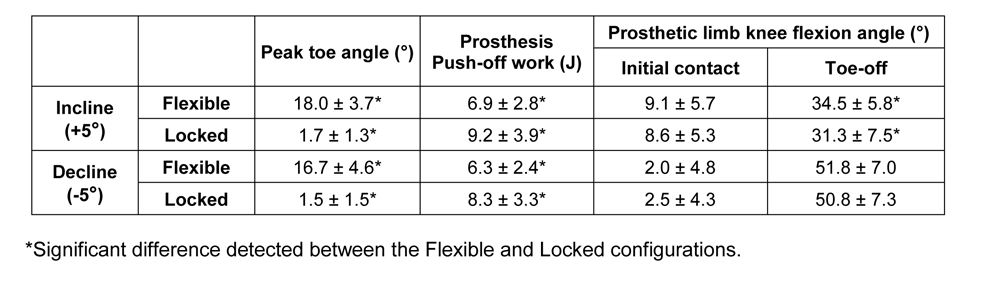
Mean (± standard deviation) peak toe joint angle, (re-dimensionalized) prosthesis Push-off work, and prosthetic limb knee angle at initial contact and toe-off.

### Prosthesis Push-off work

During both incline and decline walking, participants exhibited a decrease in Push-off work when using the Flexible toe joint versus the Locked toe joint (both p=0.008; Table 3). This was 25% (∼2 J) less prosthesis Push-off work for both incline walking (6.9 vs. 9.2 J, respectively) and decline walking (6.3 vs. 8.3 J, respectively).

### Prosthetic limb knee kinematics

For both incline and decline walking, there was no significant difference in prosthetic limb knee angle at initial contact between configurations (incline: p=0.59; decline: p=0.32; Table 3). However, during incline walking, participants had ∼3° more prosthetic limb knee flexion at toe-off for the Flexible configuration (34.5 ± 5.8°) compared to the Locked (31.3 ± 7.5°; p=0.008). During decline walking, there was no significant difference between configurations for prosthetic limb knee angle at toe-off (p=0.14; Table 3).

## Discussion

The aim of this study was to characterize the preferences and gait biomechanics of unilateral below-knee prosthesis users walking on an incline and decline wearing a prosthesis in two configurations: a Flexible and Locked out toe joint. The majority of participants (six of nine) preferred the Locked toe joint for incline and decline walking. Participants exhibited less prosthesis Push-off work during both incline and decline walking when walking with a Flexible toe joint. During incline walking, the Flexible configuration resulted in slightly more prosthetic limb knee flexion at toe-off compared to the Locked configuration.

The reduction in positive prosthesis Push-off work observed for the Flexible versus Locked configuration was statistically significant but small (incline: 6.9 vs. 9.2 J respectively; decline: 6.3 vs. 8.3 J, respectively). During incline walking at similar grades and speeds, the biological ankle/foot complex provides more than 25 J of positive power (6). Thus, the ∼2 J difference in prosthesis work between configurations may not be impactful given the 15+ J deficit in ankle work when compared to biological magnitudes. For decline walking, positive ankle/foot work is estimated to be ∼15 J (6). Therefore, a 2 J difference in prosthesis work may or may not be impactful. However, for decline walking, maximizing prosthesis Push-off work is likely not the most important determinant of an LLPU’s locomotive ability. Walking downhill does not require high levels of positive joint or center-of-mass work (Figs 4 and 5), and instead negative ankle/foot work for controlled lowering may be more important (3,6,13).

The kinematic and kinetic profiles of both intact and prosthetic limb joints were similar for the Flexible and Locked toe joint configurations, as depicted in Figs 2-5. Decreased prosthetic limb knee angle at initial contact is a common compensation employed by LLPUs to accommodate for the lack of dorsiflexion provided by typical prosthetic ankles (8,15). Prosthetic interventions that modulate ankle dynamics have had some success in improving this metric by dorsiflexing the ankle joint during swing which emulates how individuals without limb loss adapt to walking uphill (15). However, we anticipated that toe joint flexion would occur only in mid to late stance, and therefore correctly hypothesized that knee joint angle at initial contact would be unaltered by the toe joint configuration. During decline walking, increased prosthetic limb knee flexion at toe-off is considered a compensation strategy that possibly stems from the lack of ankle range of motion and stiff foot of typical prostheses (8,15). As such, we hypothesized the Flexible toe joint would provide users with additional flexibility in their ankle-foot device that could assist in lowering during late stance and therefore reduce knee flexion compared to the Locked configuration. However, we did not see a difference in prosthetic limb knee flexion at toe-off during decline walking between the Flexible and Locked configurations. It is possible that 18° of toe joint flexion at the distal end of a prosthesis does not provide adequate flexibility to compensate for the limited flexion available at the ankle during late stance. During incline walking, we did observe a 3° increase in prosthetic limb knee flexion at toe-off with the Flexible configuration. Because previous work has reported LLPUs have reduced prosthetic limb knee flexion during stance on inclines compared to controls (8,15), this increase may indicate the knee angle results in the Flexible configuration are more similar to control data. However, this difference in angle is small and the plot of average prosthetic limb knee angle throughout the gait cycle is highly similar between configurations (Fig 2).

This is the fourth manuscript in a series of studies evaluating the effects of adding a toe joint to a passive foot prosthesis during different forms of locomotion. Across all studies, we have evaluated level ground walking, sloped walking, stair ascent and decent, and uneven terrain walking with eight or nine unilateral below-knee prosthesis users (28–30). For several tasks, we observed that the Flexible toe joint reduced prosthesis Push-off work compared to the Locked toe joint. This aligns with observations of able-bodied individuals walking on level ground using a similar device where increasing toe stiffness resulted in greater prosthetic and center-of-mass Push-off work (22). Reductions in effective foot length, i.e., the anterior displacement of the center-of-pressure under the foot expressed as a percentage of total foot length (38), may be responsible. The Locked-out toe joint would enable the center-of-pressure to extend more anteriorly as it is stiffer at the distal end of the foot. In turn, the prosthetic ankle has the potential to generate larger ankle moments, and subsequently store and return more energy during walking (22,38,39). Reducing Push-off work could be considered a negative result in some instances as typical passive devices already provide reduced amounts of Push-off power compared to what is provided by the biological ankle (10,40). However, for certain tasks, maximizing Push-off work is likely not the primary factor limiting the ability of LLPUs. Tasks such as stair decent and decline walking do not require high amounts of positive joint or center-of-mass work (Figs 4 and 5) and instead controlled lowering (negative work) may be more important for LLPUs to feel comfortable and stable during locomotion.

Relatively few differences in the kinematics and kinetics of other lower limb joints were observed at the group level across the tasks tested in the full protocol. When walking on uneven terrain, a consistent reduction in prosthetic limb positive hip joint work was found (30), but most other noteworthy outcomes related to subject-specific observations. When considering the collective results from these studies, they suggest that adding an articulating toe joint likely has minimal impacts on overall biomechanics of active unilateral LLPUs. A clinician’s decision to prescribe a prosthesis with a toe joint may be best guided by individual user preference and considerations outside of the evaluated biomechanical measures (e.g., user stability).

By considering the preference results from our full protocol, the subject-specific, task-specific nature of user preference is highlighted (Table 4). User preference was mixed across the six tasks, with three to five users preferring the Flexible toe joint for each task. The Flexible toe joint was only preferred by the majority of participants during one task: walking on uneven terrain. When comparing level, incline, and decline walking, three participants had split preferences between the two configurations. Only four participants had a consistent preference for the Flexible or Locked configuration across all tasks, with the remaining participants preferring the Flexible configuration for some tasks and the Locked for others. Given that preference may be user and task specific, the selection, fitting, and alignment of a prosthetic device may benefit from evaluation across multiple tasks, beyond level ground walking. Additionally, researchers and developers should consider user preference during device design as it can be key for user acceptance (41) and note that preference for a device or behavior likely varies between tasks. Further investigation into determinants of user preference is warranted—particularly in relation to biomechanical outcomes associated with passive prosthetic devices.

**Table 4.** Participant preference for Flexible versus Locked toe joint configuration across tasks tested in the full protocol (28–30)

| Participant ID | Level Walking | Incline Walking | Decline Walking | Stair Ascent | Stair Decent | Uneven Terrain Walking |
| --- | --- | --- | --- | --- | --- | --- |
| 1 | **Flexible** | **Flexible** | **Flexible** | **Flexible** | **Flexible** | **Flexible** |
| 2 | Locked | Locked | **Flexible** | Locked | Locked | **Flexible** |
| 3 | **Flexible** | **Flexible** | **Flexible** | Locked | **Flexible** | **Flexible** |
| 4 | **Flexible** | **Flexible** | Locked | **Flexible** | Locked | **Flexible** |
| 5 | **Flexible** | Locked | Locked | **Flexible** | **Flexible** | Locked |
| 6 | Locked | Locked | Locked | Locked | Locked | Locked |
| 7 | Locked | Locked | Locked | Locked | Locked | Locked |
| 8 | Locked | Locked | Locked | Locked | Locked | Locked |
| 9 | Locked | Locked | Locked | Locked | Locked | **Flexible** |

The participants for this series of studies were all active individuals with a high level of functional ability (all K3-K4). From both local clinicians and LLPUs, we received feedback that adding a Flexible toe joint to a prothesis may be more beneficial for lower-activity individuals with unilateral limb loss or individuals with bilateral limb loss. It was proposed that a toe joint may play an important role in user stability and comfort for activities of daily living like picking up an item off the ground, turning corners, or reaching for an object on a shelf. This could be valuable for users who have limited community ambulation and desire a prosthetic ankle-foot device that provides flexibly during daily tasks. Future work could evaluate adding a Flexible toe joint to a prosthesis in a population of lower mobility users and assess tasks outside of locomotion—particularly given the challenges and potentially harmful movement adaptations that have been observed for LLPUs across a number of activities of daily living (9,42,43).

## Conclusions

The current study characterizes the effect of adding a Flexible toe joint to a passive foot prosthesis during sloped walking for active individuals with unilateral, below-knee limb loss. Three of nine participants preferred the Flexible toe joint over the Locked toe joint for incline and decline walking. We saw statistically significant changes in prosthesis Push-off work during both sloped conditions with the Flexible configuration providing ∼2 J less work than the Locked. Overall, results indicated that adding a toe joint to a passive foot prosthesis had a relatively small effect on joint kinematics and kinetics during sloped walking. This work is the fourth manuscript of a multi-part series that assessed the impact of a prosthetic toe joint across six locomotor tasks (walking over level, incline, decline, and uneven terrain surfaces, and ascending/descending stairs). The collective findings from this dataset demonstrate that user preference for passive prosthetic technology may be both highly subject-specific and task-specific. We conclude that toe joints do not appear to substantially or consistently alter lower limb mechanics for highly active (K3-K4) unilateral below-knee prosthesis users. Future work could investigate the preference and potential benefit of a prosthesis with a toe joint with lower-mobility individuals.

## Acknowledgements

The authors would like to thank Drs Gerasimos Bastas and Eric Honert for their early input on study design and prosthetists Justin Darm, Chris Arnold, and Paul Halkiades for their recruitment assistance and for performing prosthesis fittings/alignments. Additional thanks to Justin Cruz, John Kerr, Courtney Klapka, Olivia Cook, Jonathan Powles, Tristan Gilbert, and Jacob Rogatinsky for assisting in early device prototyping, data collection, and data processing.

